# Isolation and nitrogen removal characteristics of a novel aerobic denitrifying strain *Achromobacter xylosoxidans* GR7397

**DOI:** 10.1101/2023.05.24.542219

**Authors:** Ailu Gu, Yunlong Li, Wei Yao, Anqi Zhang, Zimin Chai, Maosheng Zheng

## Abstract

Aerobic denitrifying bacteria have the potential for engineering applications due to the efficient nitrate removal capacity from wastewater. In this study, a novel aerobic denitrifying strain was isolated and identified as *Achromobacter xylosoxidans* GR7397 from the activated sludge of a wastewater treatment plant, which possessed efficient nitrate removal capacity. Moreover, the denitrification capacity and properties of the strain were investigated in the presence of nitrate as the only nitrogen source. Five denitrification reductases encoding genes were harbored by strain GR7397 determined by electrophoretic analysis of PCR amplification products, consisting of periplasmic nitrate reductase (NAP), nitrate reductase (NAR), nitrite reductase (NIR), nitrous oxide reductase (NOS), and nitric oxide reductase (NOR), demonstrating that the strain has a complete denitrification metabolic pathway. The optimum denitrifying condition of strain GR7397 included sodium acetate adopted as the electron donor, COD/TN ratio at 4, pH at 8, temperature at 30°C, under which condition, the nitrate removal rate reached 14.86 mg · L^-1^ · h^-1^ that the 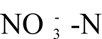 concentration decreased from 93.90 mg/L to 4.73 mg/L within 6 h with no accumulation of nitrite. In addition, the bioaugmentation performance of strain GR7397 to enhance nitrate removal was evaluated to be effective and stabilized in a sequential batch reactor (SBR). The removal rate of 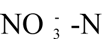 was the highest during each cycle with a range of 15.48-28.56 mg·L^-1^·h^-1^ in the SBR with inoculating 30% of the strain concentrate. The current research demonstrated that strain GR7397 has significant potential for application in enhancing nitrogen removal in wastewater treatment.

## 1. Introduction

Nitrogen is one of the most important pollutants contributing to water pollution when it exceeds a restricted concentration in the aquatic environment, which can cause deterioration and eutrophication of water quality (Zhu et al., 2012), consequently threatening the ecosystem functions of freshwater, aquaculture safety, and sanitation (Tang et al., 2019). Therefore, the development of efficient nitrogen removal technology has received widespread attention from an increasing range of researchers. Biological denitrification was gradually focused to be the most promising method to reduce total nitrogen in the water because of its safety, high efficiency, and economic efficiency. Traditional biological nitrogen removal technology utilizes nitrifying and denitrifying bacteria to convert ammonia to nitrate and then reduce nitrate to nitrogen, with the aim of achieving the removal of total nitrogen from water (Khardenavis et al., 2007). Aerobic denitrification is an emerging biological denitrification technology in recent years, which has unique advantages compared with traditional anoxic denitrification and facultative aerobic denitrification. The aerobic denitrifying bacteria could perform the denitrification step of converting nitrate to nitrite in the presence of relatively high dissolved oxygen due to the low affinity for oxygen of periplasmic nitrate reductase (NAP) (Sparacino-Watkins et al., 2014). In particular, heterotrophic nitrifying and aerobic denitrifying microorganisms have been increasingly studied since *Thiosphaera Pantotropha* was isolated in the 1980s (Lesley & Kuenen, 1983). Subsequently, other bacterial genera such as *Acinetobacter* (Su et al., 2017), *Enterobacter* (Guo et al., 2016), *Halomonas* (Ren et al., 2019), *Marinobacter* (Liu et al., 2016), *Pseudomonas* (Huang et al., 2015; Zhao et al., 2018; Zheng et al., 2014), *Fusarium* (Cheng et al., 2020), *Hanseniaspora* (Zhang et al., 2018), *Paracoccus* (Shi et al., 2013), *Vibrio* (Duan et al., 2015). *Bacillus* (Rout et al., 2017), and *Klebsiella*(Li et al., 2019), were sequentially isolated from activated sludge, industrial wastewater, rice soils, etc. With the widespread attention on aerobic denitrifying bacteria, their denitrification characteristics including influence by nutritional and physical factors, metabolic pathways, and applications in practical engineering have been investigated (Fu et al., 2022).

To date, although increasing aerobic denitrifying microorganisms have been isolated and identified, their 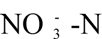 removal rates hardly meet the requirement of practical application. Two heterotrophic nitrifying aerobic denitrifying bacteria, *Achromobacter sp.* HNDS-1 and *Enterobacter sp.* HNDS-6, were isolated from paddy soil with 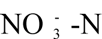 removal rate of 5.84 mg· L^-1^ · h^-1^ and 10.54 mg· L^-1^ · h^-1^, respectively (Liu et al., 2023). *Hanseniaspora uvarum* KPL 108 was isolated from drinking water reservoir sediments, which decreased 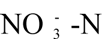 concentration from the initial 138 mg/L to 1.07 mg/L in 35 hours with an average removal rate of 3.91 mg·L^-1^·h^-1^ (Zhang et al., 2018). Similarly, it was reported that the nitrate removal rates of *Fusarium solani* RADF-77 (Cheng et al., 2020), *Rhodococcus sp.* CPZ24 (Chen et al., 2012), *Vibrio diabolicus SF16* (Duan et al., 2015) and *Marinobacter sp.* F6 (Zheng et al., 2012) reached 4.43 mg · L^-1^ · h^-1^, 3.10 mg · L^-1^ · h^-1^, 2.83 mg · L^-1^ · h^-1^, and 1.46 mg · L^-1^ · h^-1^, respectively. Accordingly, to the best of our knowledge, few strains with high-efficient aerobic denitrification properties have been reported.

For exploring aerobic denitrifying bacteria with high denitrification capacity, a novel aerobic denitrifying strain GR7397 was isolated in this research. To investigate the optimal culture environment for aerobic denitrification bacteria with a high-efficiency nitrate removal rate and to validate the feasibility of applying it to practical engineering. An aerobic denitrifying strain GR7397 with high-efficient denitrification performance was isolated and purified from the activated sludge of a wastewater treatment plant in this study. The mechanism of aerobic denitrification was explored based on the successful amplification of five aerobic denitrification functional genes. Moreover, the denitrification performance under different carbon sources, COD/TN ratios, O_2_ concentration, and pH values were also investigated. Further, the strain was inoculated into a sequencing batch reactor (SBR) for the treatment of wastewater that containing high concentrations of nitrate, and the feasibility of strain GR7397 in practical wastewater treatment applications was comprehensively evaluated through comparison of the system without bioaugmentation.

## 2. Materials and methods

### 2.1. Sludge sample and medium

Activated sludge samples were collected from the aeration tank of a wastewater treatment plant (WWTP) in Beijing, China (40°01ʹN, 116°29ʹE). This WWTP was operated with a typical anaerobic anoxic aerobic process with an HRT of 12h and SRT of 30 days (Zheng et al., 2021). Other operational parameters including influent total nitrogen (TN) concentration of 32.5 mg/L, pH of 7.3, and the dissolved oxygen of 5.2 mg/L. The mixed liquor suspended solids (MLSS) were about 8447mg/L.

The aerobic denitrifying bacterial enrichment medium (EM) consisted of (per liter) 1.71 g CH_3_COONa, 0.6 g NaNO_3_, 1.6 g K_2_HPO_4_, 0.1 g MgSO_4_・7H_2_O, 0.02 g CaCl_2_, 0.005 g FeSO_4_・7H_2_O, and 0.1 mL trace elements solution (Yao et al., 2013). The medium for screening bacteria consisted of an additional 2% agar and 1 mL of 1% BTB solution (1% bromothymol blue dissolved in anhydrous ethanol) based on EM. The aerobic denitrification ability of the strain was initially determined based on the color change by the medium as increasing pH ascribed from denitrification would make the BTB medium appear blue (Naoki et al., 2003). Denitrification medium (DM) consisted of (per liter) 0.6 g NaNO_3_, 0.512 g CH_3_COONa, 1.6 g K_2_HPO_4_, 0.1 g MgSO_4_・7H_2_O, 0.02 g CaCl_2_, 0.005 g FeSO_4_・7H_2_O, and 3 ml trace element solution. All culture media were adjusted to pH 7.5-8.0 using 0.1 M HCl and 0.1 M NaOH, and were sterilized by autoclaving at 121°C for 30 min before incubation.

### 2.2. Isolation and taxonomical identification of the strain GR7397

Erlenmeyer flask with volume of 250 mL was filled with 100 mL steriled EM and inoculated with 30 mL activated sludge sample. The inoculated flask was incubated thermostatically for 2 days on a shaker at 30°C and 130 rpm. Afterward, the cultured bacterial suspension (1 mL) was transferred to another 250 mL Erlenmeyer flask containing 100 mL EM, where the heterotrophic denitrifying bacteria were cultured at 30°C and 130 rpm for another 2 days. This inoculation culture procedure was repeated three times consecutively. Gradient dilutions of the replicate cultures were performed to obtain bacterial suspensions (multiplication of dilutions 10^-2^ - 10^-5^). 0.1 mL of bacterial suspension was taken using the plate scribing method on enriched agar solid medium and then incubated in a thermostatic chamber at 30°C. In case of visible colony formation pick a single colony onto a new enrichment agar solid medium for scribing and repeat twice to obtain purified single colonies, and were tested separately for denitrification activity in BTB solid medium. Finally strain GR7397 was investigated further as it demonstrated the most high-efficient nitrate removal rate under aerobic conditions in batch experiments. The isolated and purified aerobic denitrifying strain GR7397 was inoculated into the slant solid medium and was sent to China General Microbiological Culture Collection Center for strain conservation. Total genomic DNA was extracted from the bacterial suspension using the Bacterial DNA Isolation Kit (Chengdu Fuji Biotechnology Co., Ltd.) according to the manufacturer’s instructions. The gene encoding 16S rDNA was amplified from genomic DNA by polymerase chain reaction using forward primer F27 (5’-AGAGTTTGATCATGGCTCAG-3’) and reverse primer R1492 (5’-TACGGTTACCTTGTTACGACTT-3’) (Shanghai Shengong Bioengineering Company). The PCR amplification reaction conditions were as follows: 94 °C for 3 min; 24 cycles, 94 °C for 30 s; 54 °C for 30 s; 72 °C for 1.5 min, and extended at 72°C for 10 min(Zheng et al., 2014). PCR products sequenced by the Sanger method. The sequencing results were analyzed using BLAST software (BLAST: Basic Local Alignment Search Tool (nih.gov)), and the spliced sequences were compared with other known sequences in the NCBI database. Simultaneously, a phylogenetic tree was constructed based on the Neighbor-joining method using MEGA 7. 0 software.

### 2.3. PCR amplification of denitrification functional genes

5 mL purified bacterial suspension was obtained for the analysis of denitrification functional genes. Total genomic DNA in bacterial suspensions was extracted from the sample using the FastDNA Spin Kit (MP Biomedical, Illkirch, France) according to the manufacturer’s instructions, and stored at −20 °C until use. DNA concentration and purity were determined with a microspectrophotometer. Denitrification genes including periplasmic nitrate reductase gene (*napA*), nitrate reductase gene (*narG*), nitrite reductase gene (*nirS*), nitrous oxide reductase gene (*nosZ*), and nitric oxide reductase gene (*cnorB*), were determined by q-PCR technique according to the primers shown in Table 1. Operating conditions on Applied Biosystem 7300 were as follows: pre-denaturation at 94 °C for 15 min; 35 cycles of denaturation at 95 °C for 30 s, annealing for 1 min at 58 °C for *narG* and *napA*, at 60 °C for *nirS*, at 56 °C for *cnorB* and at 60 °C for *nosZ*, extension at 72 °C for 45 s for *napA* and *nirS*, or 60 s for *cnorB* and *nosZ*; the final step was an extension at 72 °C for 1 min. The total volume of PCR amplification was 20 μL, containing 2 μL DNA template, 15 μL PCR ReadyMix (BGI, Shenzhen, China), 0.5 μL of each primer (10 μmol/L) and 2 μL DD H_2_O. DD H_2_O was selected to replace the DNA template for a blank control to exclude external interference. The amplified PCR products were visualized by electrophoresis on a 1.8% agarose gel stained with ethidium bromide.

**Table. 1.**
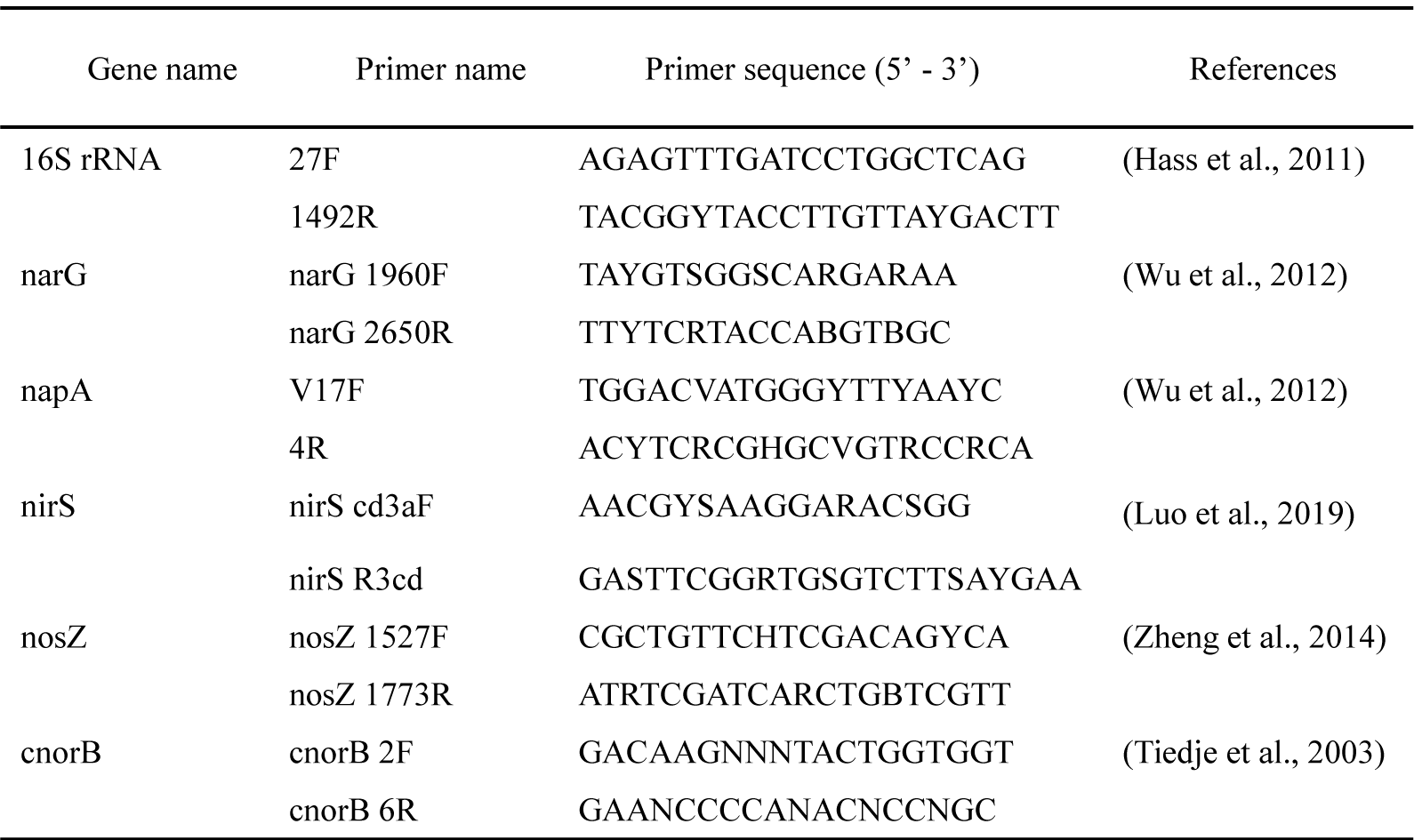
PCR primers used for 16S rRNA gene sequencing and qPCR analysis.

### 2.4. Optimal culture conditions for aerobic denitrification of strain GR7397

To investigate the maximum denitrification rate of aerobic denitrifying strain GR7397, different environmental factors that would promote or inhibit aerobic denitrification, which included carbon source supply, COD/TN ratio, pH, and oxygen concentration were investigated. The condition which optimized the denitrification capacity of strain GR7397 would be taken as the condition in the next factor experiment. The effects of methanol, sodium acetate, sodium citrate, sodium succinate, glucose, and ethanol were selected as various carbon sources for the aerobic denitrification capacity of strain GR7397. Subsequently, the COD/TN ratio was selected as 2, 4, 6, 8, and 12 in the culture medium. When investigating the effect of COD/TN, the 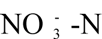 concentration in the culture medium was kept constant at 100 mg/L, with changing the concentration of the carbon source to vary the different COD/TN ratios at 2, 4, 6, 8, and12, respectively. Furthermore, oxygen concentration influence on the denitrifying strain was investigated. The anaerobic flask was filled with helium for 15 min to evacuate the headspace and then charged with a relative proportion of oxygen so that the relative proportion of oxygen in the flask were 0%, 10%, 20%, 30%, 50%, and 100%, respectively. Eventually, while exploring the effect of pH on denitrifying strain GR7397, the pH of the medium was adjusted to 6±0.2, 7±0.2, 8±0.2, 9±0.2 and 10±0.2 by 0.1 M sodium hydroxide or 0.1 M dilute hydrochloric acid.

Before each batch of experiments, strain GR7397 was activated for culture. 5 mL of the purified strain concentrate was injected into a 250 mL Erlenmeyer flask containing 100 mL of DM with a sterile syringe and incubated in a thermostatic shaker at 30°C and 130 rpm until the OD>0.8 of the culture. Thereafter, 40 mL of DM was added to the 100 mL anaerobic flask corresponding to the various influencing factors. 2 mL of the activated bacterial solution was inoculated into the anaerobic bottle, which was incubated in a shaker at 30°C and 130 rpm, with the concentration of nitrate-nitrogen and nitrite-nitrogen measured every 2 h until the nitrate nitrogen was basically consumed. A series of parallel experiments were set up for each group to exclude the effect of errors on the experimental results.

### 2.5. Potential applications of strain GR7397 in activated sludge systems

To investigate the feasibility of the isolated denitrifying strain GR7397 for application in the wastewater treatment. A serum bottle with a volume of 1L was selected as the sequencing batch reactor (SBR). Different amount of strain GR7397 concentrate (0%, 15%, and 30%) and a certain volume of activated sludge were inoculated the SBRs with working volume of 300 mL (MLSS of 2000-2500 mg/L). In particular, the activated sludge was domesticated and cultivated in the aeration tank of the reactor for two sludge retention time at hydraulic retention time (HRT) of 12 h, a 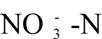 concentration of 100 mg/L, and COD/TN ratio of 4. The serum flasks were sequentially supplemented with DM that satisfied the optimal culture conditions for strain GR7397 and 30% hydrophilic polyurethane foam (1 m³, Qingdao Banghao Co., Ltd.) (Feng et al., 2012) to ensure a final effective liquid volume of approximately 300 mL. SBRs were incubated in a thermostatic shaker at 30°C and 130 rpm. One cycle of SBRs operation was established for 12 h, including 5 min of feeding, 11 h of operating, 50 min of settling, and 5 min of decanting. The hydrophilic polyurethane packing with adsorbed activated sludge and GR7397 strain was retained after 50 min of decanting and 200 mL of supernatant was withdrawn using a sterile syringe. Thereafter, the serum bottle was filled with 200 mL of denitrifying concentrated liquid medium to ensure that the effective volume in the SBR was 300 mL and that the content of the various substances inside the bottle was the same as the DM after shaking uniformly. The concentrations of 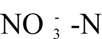 and 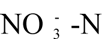 were determined from the second cycle at an interval of hour in SBRs. A set of parallel experiments were set up for each group.

### 2.6. Analytical methods and calculations

The culture fluids were separated by centrifugation at 12,000 rpm in the high-speed benchtop microcentrifuge with the supernatant for chemical analysis after centrifugation. 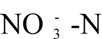 or 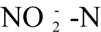 was determined using SmartChem 200 fully automated chemistry analytical instrument (AMS Alliance, Italy).

The removal efficiency (%) and rate mg·L^-1^·h^-1^ of 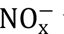 was calculated as follows:

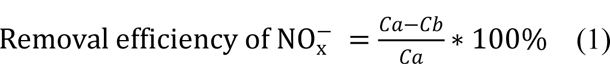

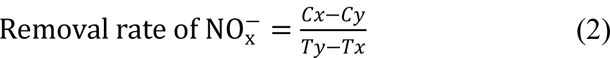

*Ca* and *Cb* (mg/L) are the concentrations of 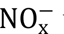 at the beginning and end of the experiment, respectively; *Tx* is the start time and *Ty* is the end time of the experiment, *Cx* and *Cy* (mg/L) correspond to the concentrations of 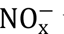 at *Tx* and *Ty*, respectively (Ke et al., 2022).

## 3. Results

### 3.1. Identification of the strain GR7397

After serial enrichment and purification, an aerobic denitrifying strain with high-efficiency denitrification ability was isolated from the activated sludge of a WWTP, which was named as GR7397. The phylogenetic tree including this strain and other adjacent strains was constructed using 16S rDNA sequencing results compared with the GenBank database (Fig. 1). The results indicated that the strain belonged to the genus *Achromobacter* and was 99.93% similar to *Achromobacter xylosoxidans* Hugh 2838. Additionally, it should be mentioned that the 16S rDNA gene sequence of *Achromobacter sp.* GR7397 was not identified in the NCBI nucleotide database, demonstrating that this strain was screened for the first time and identified as *Achromobacter xylosoxidans* GR7397.

**Fig. 1.**
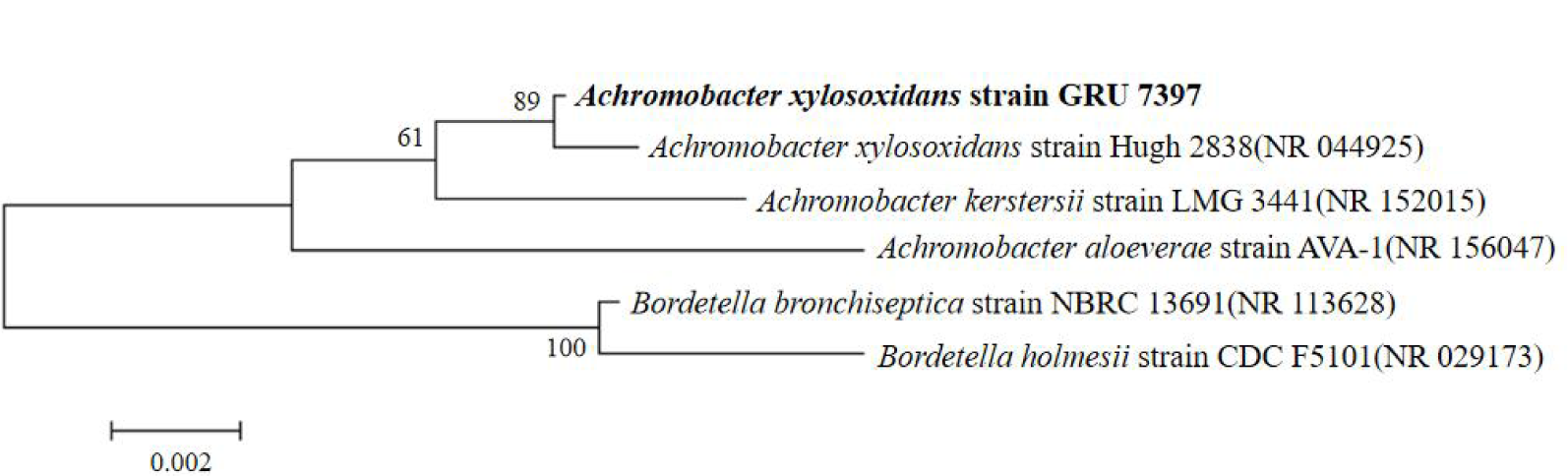
Phylogenetic tree drawn from neighbor-joining analysis based on the 16S rDNA gene sequence of *Achromobacter xylosoxidans* strain GR7397 and other reference sequences.

### 3.2. PCR amplification of denitrifying enzyme genes

PCR amplification of genes related to aerobic denitrification was performed to study the nitrogen metabolism pathway of strain GR7397. Five potential nitrogen metabolizing denitrifying enzyme encoding genes were examined by agarose gel electrophoresis using PCR amplification products containing, *narG*, *napA*, *nirS*, *cnorB* and *nosZ*. The stained electrophoresis gel plates were visualized under a long-wave UV illuminator at a wavelength of 245 nm. As the results shown in Figure 2, it is evident that strain GR7397 denitrifying genes (Fig. 2). Thereby it could perform the complete metabolic pathway, i.e., 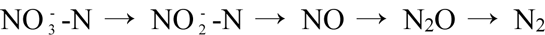. The corresponding reductase is required for each step of the reduction process. Microbial nitrate reductases are classified as allosteric nitrate reductase (NAR), periplasmic nitrate reductase (NAP), and anabolic nitrate reductase (NAS) based on the physiological function of the enzyme, the subcellular location, and the structure of the molybdenum active sites. (Sparacino-Watkins et al., 2014). The enzyme that catalyzes the reaction of reducing the nitrous oxide to nitrogen gas is nitrous oxide reductase (NOS) in microorganisms, encoded by the *nosZ* gene, which is responsible for the final step of the denitrification process (Roy et al., 2021). Nitrous oxide in most denitrifying microorganisms, like the present strain GR7397 is an intermediate during the denitrification process, while nitrous oxide in some microbes laking nosZ gene is the terminal product, which would pose an considerable emission of the unwanted greenhouse gas (Kuypers et al., 2018). As Achromobacter sp. GR 7397 contains NAR, NAP, NIR, NOR, and NOS, it demonstrated the potential to perform the complete denitrification process, realizing not only nitrate reduction but also the control of N2O emission.

**Fig. 2.**
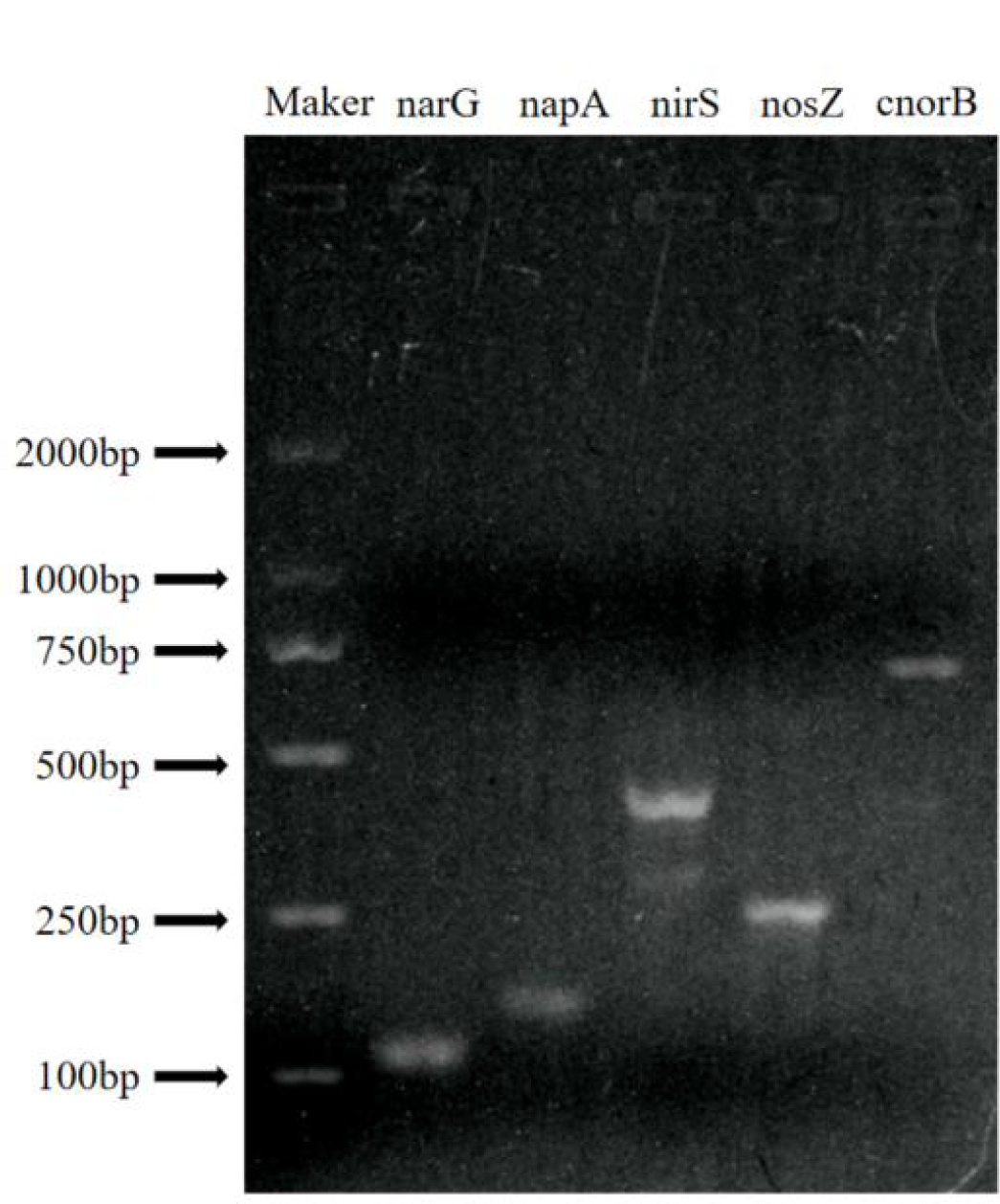
Electrophoretic analysis of PCR amplification products for *narG*, *nap*A, *nirS, cnorB* and *nosZ*.

### 3.3. Optimum culture conditions for strain GR7397

#### 3.3.1 Effect of different carbon sources on strain GR7397

In this study, methanol, sodium acetate, sodium citrate, sodium succinate, glucose, and ethanol were selected as carbon sources with nitrate as the sole nitrogen source to investigate the effects of different carbon source on aerobic denitrifying strain GR7397. The variation curves of 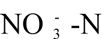 and 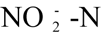 concentrations using different carbon sources to treat simulated wastewater with high nitrate nitrogen were shown in Fig. 3a. During the experiments with glucose and methanol as the only carbon source, the concentration of 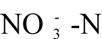 was eventually dropped from 100 mg/L to 47.60 mg/L and 65.30 mg/L over 12 h. The 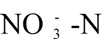 removal efficiency was 50.16% and 28.79%, respectively (Fig.3a). The reaults indicated that the methanol and glucose as carbon sources were not favorable to the cultivation of strain GR7397 when compared with other carbon sources, which significantly inhibited the denitrification ability, and delayed the initiation of the denitrification of strain GR7397. However, glucose promoted the denitrification of strain GR7397 more effectively than methanol. In the experiments with sodium acetate and sodium succinate as the carbon sources, nitrate-nitrogen was almost completely degraded at the 8th and 10th hour, while at the same time nitrite-nitrogen did not accumulate. Meanwhile, the removal rates of nitrate by sodium acetate and sodium succinate were both rapid at 6 h-8 h, reaching an average of 12.05 mg· L^-1^ · h^-1^ and 9.58 mg· L^-1^ · h^-1^, respectively. In addition, sodium acetate was the carbon source with the highest nitrate removal capacity, with a removal efficiency of 98.50% after 8 hours of nitrate denitrification; the next removal efficiency was 72.76% using sodium succinate as the carbon source. In the microcosm with sodium acetate and sodium succinate as carbon sources, almost all of the nitrate and nitrite have eventually completed their denitrification reduction reactions. In addition, sodium citrate and ethanol also could significantly promote the denitrification of GR7397, with 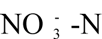 removal efficiencies of 93.71% and 93.20%, respectively. To sum up, with nitrate as the only nitrogen source and sodium acetate as the carbon source, strain GR7397 had the strongest denitrification capability and the fastest removal rate of nitrate. The denitrification capacity of strains GR7397 using organic substances as carbon sources ranged from sodium succinate, sodium citrate, ethanol, glucose, and methanol. Hence, sodium acetate was used as the carbon source in DM for the following experiments investigating the different influencing factors on the optimal culture conditions of strain GR7397.

**Fig. 3.**
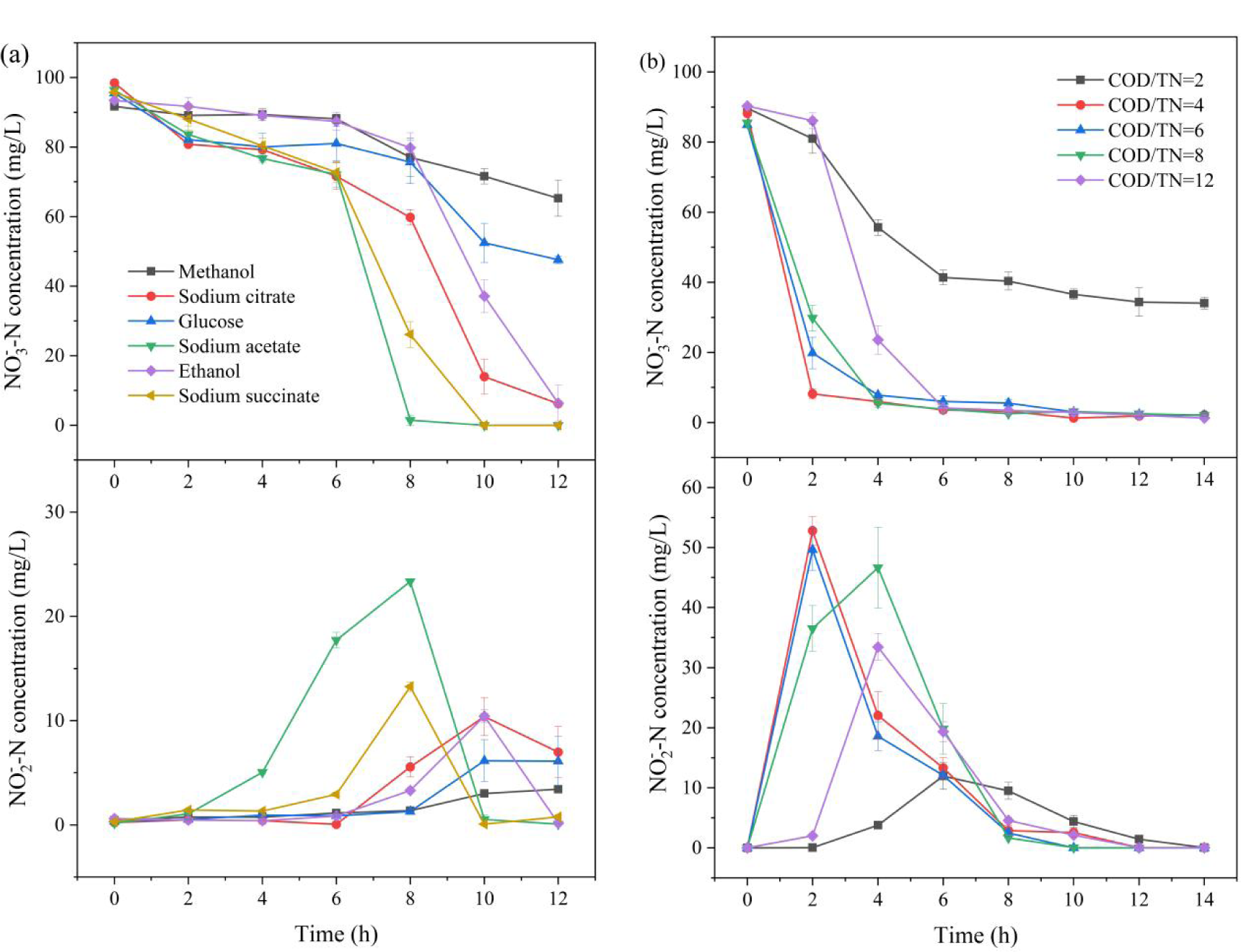
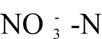 and 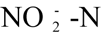 concentrations as a function of time in simulated high nitrate nitrogen wastewater inoculated with strain GR7397 with different carbon sources (a) and different COD/TN ratios (b).

#### 3.3.2 Impact of different COD/TN ratios on strain GR7397

The carbon source would provide electrons for the denitrification and the necessary energy for the aerobic respiration of microorganisms. Substrate utilization rate by aerobic denitrifying microorganisms is influenced by the ratio of carbon and nitrogen sources. Generally, overhigh COD/TN ratios would inhibit the activity of denitrifying bacteria, which leads to ineffective denitrification and leaves excess organic matter unused and wasted at the end of the reaction, while the lower COD/TN ratio is also detrimental to the denitrification reaction since there are not enough nutrients to satisfy the normal denitrification process resulting in poor denitrification (Deng et al., 2020).

In this study, the effect of the COD/TN ratio on strain GR7397 was investigated by keeping the concentration of nitrate concentration in the medium constant and changing the concentration of carbon source to achieve different COD/TN ratios, and the experimental results are shown in Figure 3b. Most of the 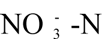 concentrations were effectively reduced at COD/TN ratios of 4-12, while 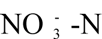 removal was discontinued at 62.04% at a COD/TN ratio of 2. However, the nitrate removal efficiency reached 90.75% by the 2nd hour when the C/N ratio was 4, which was the fastest nitrate removal rate among all COD/TN ratios. The second highest nitrate removal rate was at COD/TN ratio of 6 when the nitrate removal rate was 76.68% and the removal rate was up to 90.81% at 4 h. Although the accumulation of 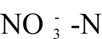 was observed in the 2nd hour at COD/TN ratios of 4 and 6, the accumulated 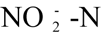 was rapidly degraded as the denitrification process advanced (Fig. 3b). The denitrification of strain GR7397 was relatively delayed, and the nitrate removal efficiency was high after the initiation of denitrification at a COD/TN ratio of 12. These results demonstrated that providing the strain with an appropriate carbon source could promote both the reduction of 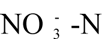 concentration and the accumulation of 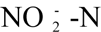, while too low or excessive carbon sources could inhibit the denitrification ability of the strain. Therefore, the COD/TN ratio of 4 with the highest removal efficiency and the fastest removal rate was selected as one of the optimal culture conditions for strain GR7397. Based on the effects of two environmental factors, including carbon source and COD/TN on the denitrification ability of strain GR7397, it was selected that sodium acetate as carbon source and COD/TN ratio of 4 in the following experiments to further explore the optimal culture conditions of the strain.

#### 3.3.3 Different oxygen concentrations

To investigate the effects of different oxygen concentrations on the growth and denitrification capacity of the facultative denitrifying bacterium GR7397, the headspace of the anaerobic flasks was filled with helium and then filled with a relative proportion of oxygen. When the initial oxygen concentration in the serum bottle was 0% and 10%, the concentration of 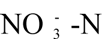 began to decline first, and the rate of nitrate removal at the 6th hour was the highest compared to other oxygen concentrations, respectively, and the removal efficiency was 89.41% and 69.77%, while the accumulation of 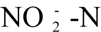 reached a peak at this time but was quickly degraded by denitrifying bacteria (Fig. 4a). The removal rate of nitrate nitrogen was 11.87 mg·L^-1^ · h^-1^ when all nitrate was reduced; denitrification rates were essentially the same at 30%, 50%, and 100% oxygen concentrations, and strain GR7397 consumed all the nitrate and nitrite in the bottle after the tenth hour at different oxygen concentrations. The denitrification capacity of strain GR7397 was not altered with oxygen concentration at oxygen concentrations above air saturation (21%). In summary, the denitrification capacity of strain GR7397 reached equilibrium at oxygen concentrations above air saturation (21%), and the denitrification capacity of the strain did not significantly differ in serum flasks with different oxygen concentrations. Hence, the denitrification capacity of strain GR7397 was not significantly affected by the oxygen content, so the oxygen concentration was not constrained in the next experiments (Fig.4a).

**Fig. 4.**
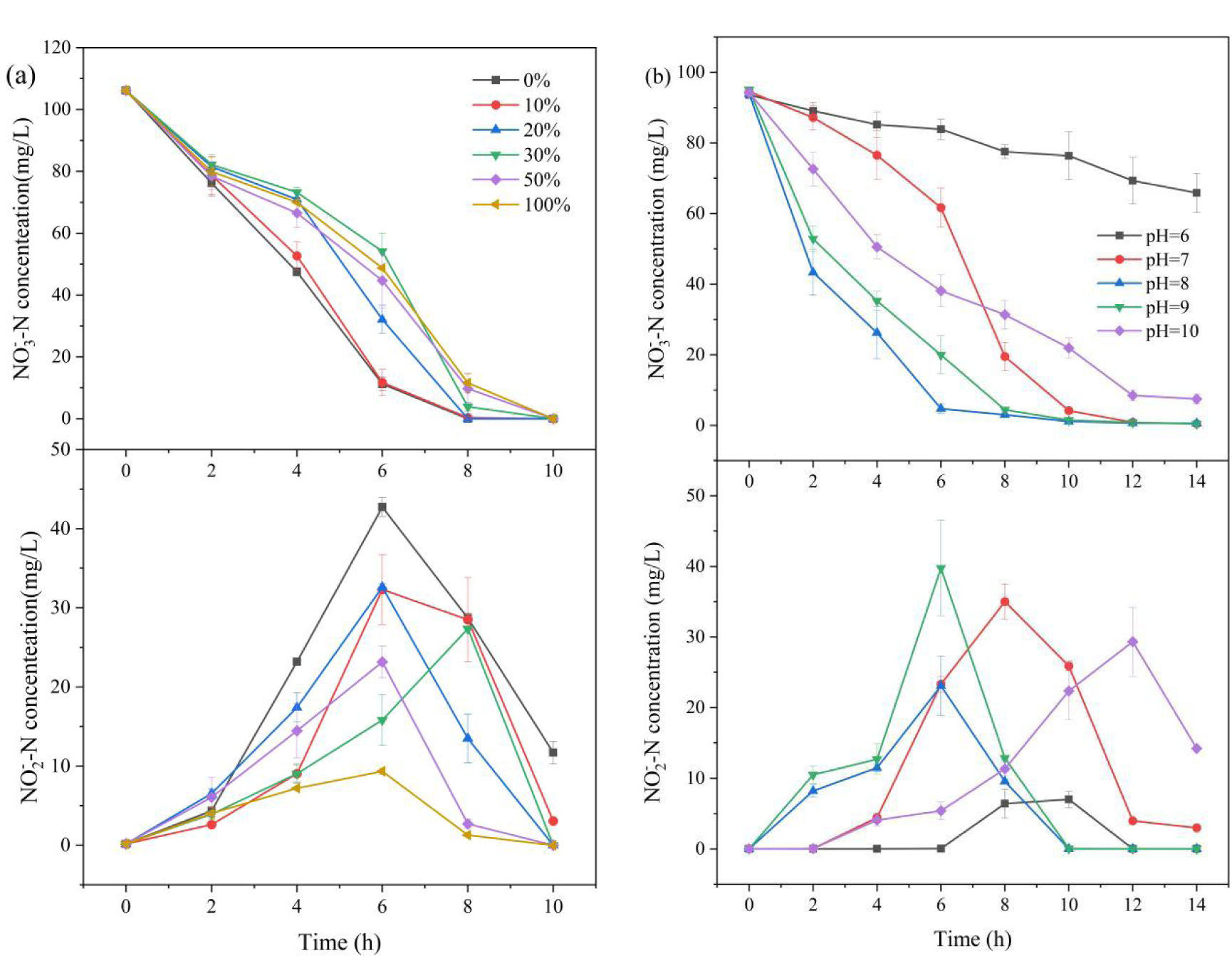
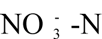 and 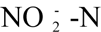 concentrations with time-dependent changes in simulated wastewater treated with strain GR7397 at different relative oxygen concentrations (a) and different pH levels (b).

#### 3.3.4 Effect of different initial pH values on GR7397

The pH of the environment is significant in the metabolic activity of microorganisms, which is the primary role being to cause changes in the charge of the cell membrane. It affects the absorption of nutrients by microorganisms, the activity of enzymes in the metabolic process, the availability of nutrients in the growth environment and the toxicity of harmful substances (Hu et al., 2021). The effects of weakly acidic and weakly alkaline environments on the denitrification performance of strain GR7397 are shown in Figure 4b. Overall, strain GR7397 exhibited stronger denitrification performance in weakly alkaline and neutral environments compared to weakly acidic conditions. The fastest nitrate removal rate of strain GR7397 occurred in pH 8, with a nitrate removal efficiency of 94.96% at the 4th hour, and the removal rate of 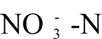 was 14.86 mg·L^-1^·h^-1^. The efficiency of nitrate removal by the strain in the microcosm reached 95.37% and the removal rate of 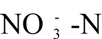 was 11.34 mg·L^-1^·h^-1^ at pH 9, which was second only to the efficiency at pH 8 when 6 hours were consumed. Meanwhile, the accumulation of 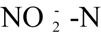 simultaneously peaked at pH 8 and 9 at the 6th hour, and the accumulation of 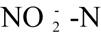 was relatively more at pH 9. The removal rate of 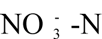 from the microcosm was slow when the pH was 6, eventually reaching a removal efficiency of 29.65% and accumulation of nitrite was not observed. The denitrification rate of the strain was weak in the first 6 hours at pH 7. After that, denitrification capacity in the microcosm increased immediately, and the accumulation of 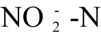 peaked at the 8th hour. At the end of the reaction at pH 7, both nitrate and nitrite nitrogen were completely degraded (Fig. 4b). As a consequence, the denitrifying strain GR7397 is most suitable for growth and reproduction in a weak alkaline environment and the strain has the strongest denitrification capacity at pH 8.

To summarize, the optimal culture conditions for strain GR7397 were when sodium acetate was used as the electron donor, the COD/TN ratio was 4, the pH was 8, and the temperature was 30°C. The best 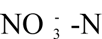 removal rate in microcosm was 14.86 mg·L^-1^·h^-1^ with 94.97% removal efficiency, and it took six hours to reduce the nitrate-nitrogen concentration from 93.90 mg/L to 4.73 mg/L with no nitrite accumulation.

### 3.4. Potential applications of strain GR7397 in activated sludge systems

According to the above study, strain GR7397 was observed to have high-efficiency nitrate removal ability and was further studied for its feasibility in practical engineering applications. Based on the experimental results in section 3.3, the following experimental conditions were formulated as that no changes were made to the oxygen concentration, sodium acetate was selected as the only carbon source, the COD/TN ratio was 4, and the pH was adjusted between 7.5 and 8.5.

Fig. 5 describes the concentration variation of nitrate and nitrite in the concentrates of strains inoculated with different concentrations in SBRs. It is significantly evident from Fig. 5a that the 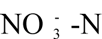 removal rate was faster in the reactor inoculated with denitrifying strain GR7397 concentrate than in the reactor without strain GR7397 concentrate, confirming that strain GR7397 had a stronger denitrification capacity when put into the SBR and that the higher inoculation amount of strain GR7397 cause the faster 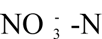 removal rate in the reactor. The relative abundance of inoculated strain GR7397 was proportional to the removal rate of 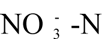. The 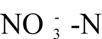 removal rates in the SBR concentrated with the denitrifying strain in the second cycle were higher than that without the strain GR7397. At the first hour, the removal rates of 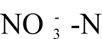 were 6.71 mg·L^-1^·h^-1^, 20.02 mg·L^-1^·h^-1^, and 29.47 mg·L^-1^·h^-1^ in the SBR without strain concentrate, with 15% strain concentrate and 30% strain concentrate, respectively. The 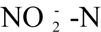 accumulated during denitrification was completely reduced in the process over eight hours. Therefore, the efficient nitrate removal capacity of strain GR7397 was significantly demonstrated in the practical water treatment process. Meanwhile, the nitrate removal rates were 12.78, 13.69, and 15.90 mg·L^-1^ ·h^-1^ in the SBR without strain concentrate, with 15% strain concentrate, and with 30% strain concentrate when the 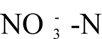 concentration was <10 mg/L, respectively. Nevertheless, 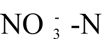 removal rates were gradually approached in the two inoculated groups of reactors after 5 cycles (Fig. 5). Such a phenomenon revealed that the increasing bacterial biomass eventually converges to an equilibrium state after the strain GR7397 grew and formed biofilm on the carrier of the SBR. Also, as can be seen in Fig. 5b, nitrite is preferentially accumulated in the reactor inoculated with 30% strain concentrate and is predominately reduced to nitrogen in each cycle. It indicates that the initial inoculum concentration would shorten the initiation process of denitrification and preferentially complete the denitrification process.

**Fig. 5.**
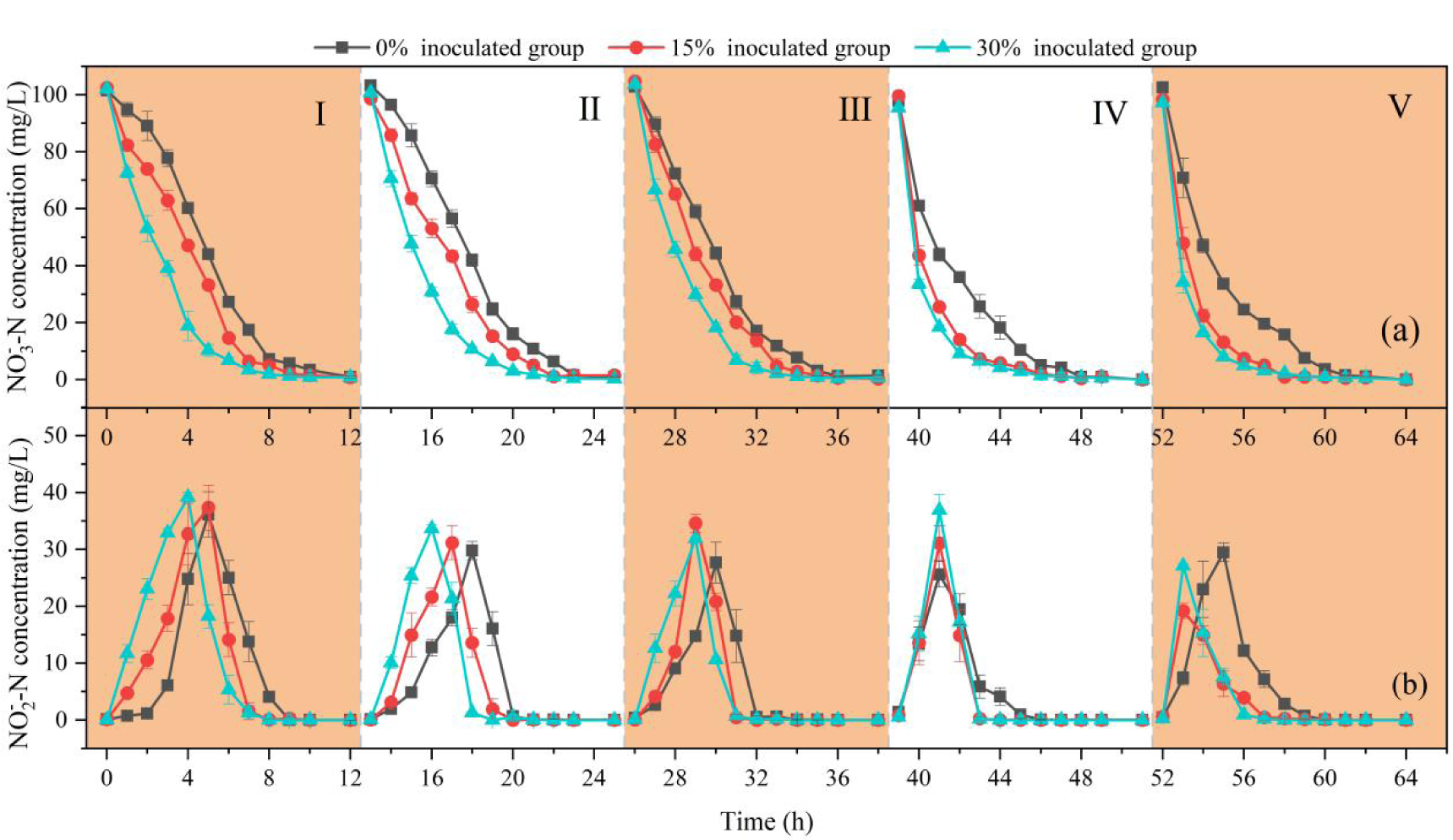
Variations of nitrate and nitrite concentrations in SBRs inoculated with different amount of strain GR7393 .

The removal rates of 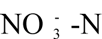 for each cycle of continuous SBR operation are depicted in Fig. 6. The 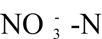 removal rate of 30% inoculated group was the highest in each cycle, with nitrate removal rates ranging from 15.48-28.55 mg · L^-1^ · h^-1^. The second highest was 15% inoculated group which removed nitrate at a rate in the range of 13.14-21.67 mg·L^-1^ ·h^-1^. The lowest nitrate removal rate was 0% inoculated group which ranged from 11.95-13.77 mg· L^-1^ · h^-1^. Overall, the 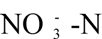 removal rate in the second cycle was significantly lower compared to the first cycle in both inoculated and non-inoculated SBRs, and accelerated rapidly from the third cycle onwards. To sum up, denitrifying bacteria were immobilized on the carrier and multiplied rapidly, which resulted in a rapid increase of nitrate removal rate. Denitrifying strain GR7397 can significantly enhance the denitrification rate, with high-efficient nitrate-nitrogen removal ability, which has the potential to be applied in the reactor.

**Fig. 6.**
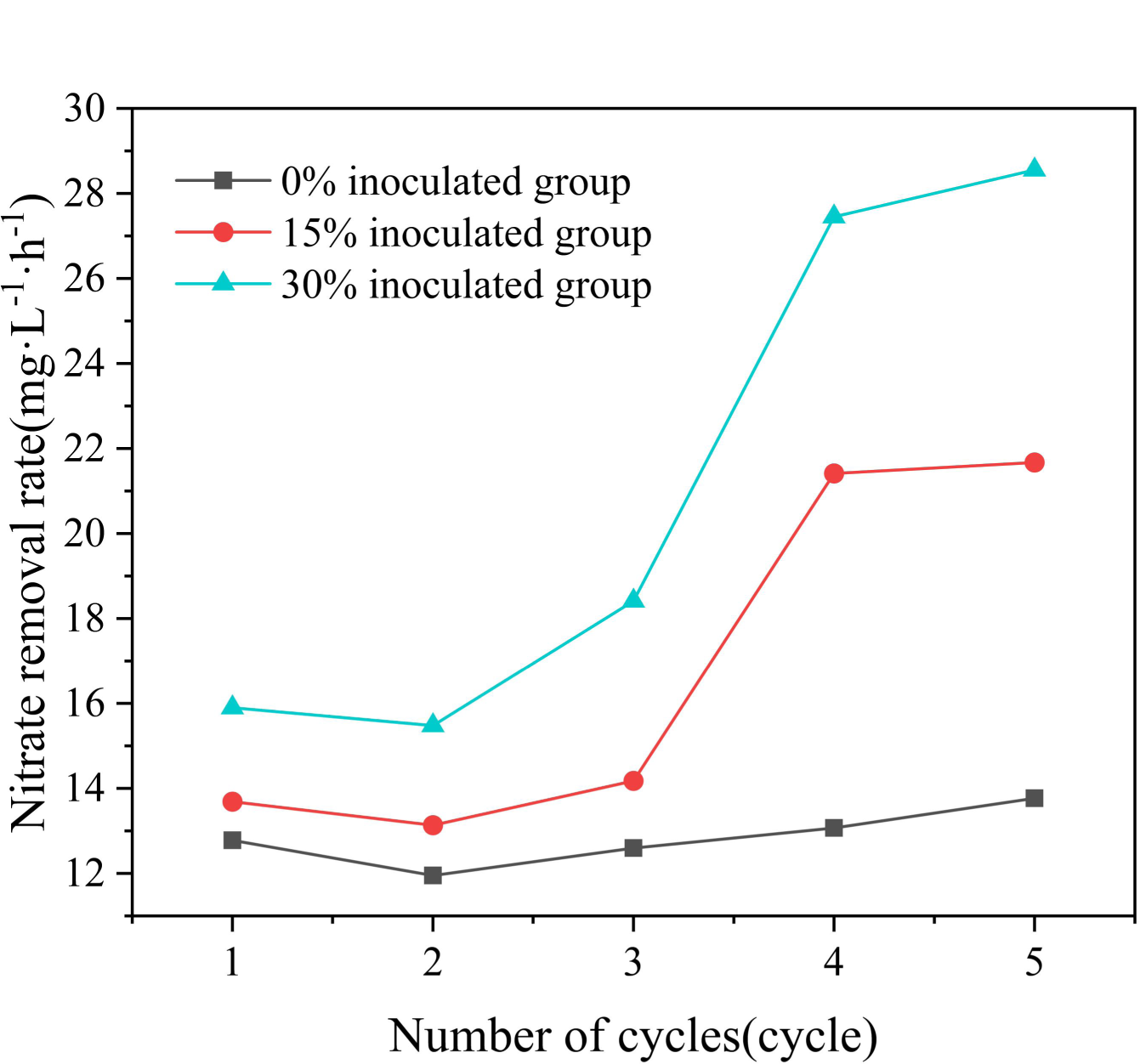
Removal rate of nitrate nitrogen in each cycle of the continuous SBR operation.

## 4. Discussion

Although aerobic denitrifying bacteria have been widely reported (Chen et al., 2020; Lang et al., 2020; Ren et al., 2019; Wang et al., 2018), most aerobic denitrifying bacteria reported in the current study denitrify at relatively lower rates. Nonetheless, the strain *A. xylosoxidans* GR7397, which was isolated from activated sludge in an aeration tank treating domestic wastewater, could have efficient performance in removing nitrate under aerobic conditions for this research. Five potential nitrogen metabolizing denitrifying enzyme genes were examined by agarose gel electrophoresis using PCR amplification products which contained *narG*, *napA*, *nirS*, *cnorB*, and *nosZ*. Strain GR7397 includes two nitrate reductases, which are NAR and NAP, nitrite reductase (NIR), nitrous oxide reductase (NOS), and nitric oxide reductase (NOR), demonstrating that strain GR7397 can complete the complete denitrification process independently. Interestingly, the presence of both NAP and NAR genes is essential to ensure that strain GR7397 could reduce nitrate to nitrite in both aerobic and anoxic environments or even in the absence of oxygen. Because NAP is insensitive to oxygen, it can be expressed under aerobic conditions, while the presence of oxygen inhibits the activity of NAR (Sparacino-Watkins et al., 2014). In parallel, the presence of NOS reductase allowed strain GR7397 to reduce nitrous oxide accumulation during denitrification, thus reducing greenhouse gas emissions (Roy et al., 2021). Hence, the strain GR7397 could be preferentially selected for application in the actual wastewater treatment process.

In this study, the effects of different carbon sources, COD/TN ratios, oxygen concentrations, and pH values on strain GR7397 were investigated for the purpose of gaining a clear insight into the growth characteristics and optimal growth environment of strain GR7397. The results showed that strain GR7397 could achieve the best denitrification treatment of wastewater with 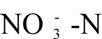 removal rate of 14.86 mg·L^-1^ ·h^-1^ when sodium acetate was used as the electron donor, COD/TN ratio was 4, pH was 8, the temperature was 30℃, and dissolved oxygen was not controlled. Due to its simple molecular structure, sodium acetate is directly utilized by GR7397 for metabolism without modification. However, methanol and glucose did not significantly promote denitrification by GR7397 because methanol must be enzymatically converted to formic acid before being metabolized by denitrifying bacteria. As well, glucose needs to be broken down into pyruvate via the glycolytic pathway before being further converted into simple alcohols and acids. The mechanism of sodium acetate utilization by strain GR7397 was more readily available compared to methanol and glucose (Jiang et al., 2022; Xu et al., 2018). Simultaneously, perhaps due to a preference for weakly alkaline environments, strain GR7397 prefers organic acids (acetate and succinate) as carbon sources rather than sugars (glucose). The reason is that when sugars are used as a carbon source, the pH after metabolism decreases due to their oxidation-reduction potential of them own (Song et al., 2021). Alternatively, the lower denitrification performance of strain GR7397 at COD/TN < 4 could be attributed to the number of electrons provided being too low to provide sufficient energy for cell growth containing a low concentration of carbon source (Kim et al., 2008). At the same time, the excessive COD/TN ratio caused a large number of other heterotrophic bacteria to multiply, resulting in competition between limited substrates, thus inhibiting the activity of denitrifying bacteria and making denitrification less effective (Deng et al., 2020). In this study, the initiation time of the denitrification capacity of the strain was prolonged by 2 hours when COD/TN=12, confirmed that the denitrification capacity of the bacteria was inhibited by an excessive carbon source. The strain GR7397 was therefore more capable of denitrification when the COD/TN ratio was between 6 and 8. The effect of pH on the rate of denitrification was mainly reflected by affecting the activity of the denitrifying enzymes. The enzyme activity would be highest in the optimum pH range, and either lower or higher pH would affect the enzyme activity. The microorganisms were died when the maximum pH was exceeded or when it was below the minimum pH (Ke et al., 2022). The nitrate removal efficiency of strain GR7397 ended up being 29.65% at pH 6. This phenomenon was probably caused by the inappropriate growth and reproduction of strain GR7397 in mildly acidic conditions and affected the denitrification performance of the strain. The most suitable pH range for most aerobic denitrifying bacteria is 7.5-8, such as *Penicillium sp.* M25-22 (Wang & Yu, 2010), *Stenotrophomonas sp.* MSNA-1 (Zeng et al., 2020), *Acinetobacter junii* YB (Prieur, 2000), *Marinobacter sp.* NNA5 (Liu et al., 2016), *Marinobacter sp.* F6 (Zhang et al., 2012), and *Bacillus sp.* N31 (Huang et al., 2017), which were consistent with the findings of strain GR7397.

This study further evaluated that the nitrate removal capacity of strain GR7397 in SBR was effective and stable. The 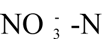 removal rate in the 30% inoculated group was the highest in each cycle, which was ranging from 15.48-28.55 mg · L^-1^ · h^-1^. Meanwhile, the removal efficiency was as high as 99% to 100%. This result is almost consistent with the performance of the SBR inoculated with Enterobacter sp. strain FL separately, with influent nitrate-nitrogen concentrations ranging from 100 −110 mg/L, setting for 24 h each cycle, with nitrate removal efficiencies ranging from 90.20 to 99.70% (Wang et al., 2018). The feasibility of strain GR7397 for wastewater treatment in open systems was further confirmed as strain GR7397 indicated significant aerobic denitrification capability in the SBR operation after mixing with microbial populations, even under uncontrolled oxygen conditions. To sum up, aerobic denitrification of this strain has great potential for application in bioreactors. Since mixed microbial population systems are typically used in WWTPs, it remains to be explored whether aerobic denitrifying strain GR7397 can be the dominant population in a sophisticated open system for wastewater treatment.

## 5. Conclusion

In this study, a novel and highly efficient aerobic denitrifying strain *Achromobacter xylosoxidans* GR7397 was isolated from the activated sludge of an domestic wastewater treatment plant to address several environmental pollution problems caused by excessive nitrogen in water. The presence of functional genes including *narG*, *napA*, *nirS*, *cnorB*, and *nosZ* demonstrated by electrophoretic analysis of PCR amplification products indicated that the strain had a complete denitrification capacity, i.e., 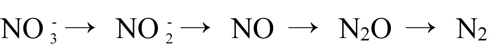. The optimum denitrification conditions for the strain were obtained with sodium acetate as an electron donor, COD/TN ratio of 4, pH of 8, dissolved oxygen not controlled. The nitrate removal rate of inoculated strain GR7397 was demonstrated to be higher than the nitrate removal rate in activated sludge systems operating in SBR. These studies reveal that strain GR7397 could be preferentially selected for the treatment of wastewater with high concentrations of nitrate or nitrite.

